# A Deep Learning Approach for Mental Health Quality Prediction Using Functional Network Connectivity and Assessment Data

**DOI:** 10.1101/2023.06.01.543257

**Authors:** Meenu Ajith, Dawn M. Aycock, Erin B. Tone, Jingyu Liu, Maria B. Misiura, Rebecca Ellis, Tricia Zawacki King, Vonetta M. Dotson, Vince Calhoun

**Author notes:** Contributing authors.

## Abstract

While one can characterize mental health using questionnaires, such tools do not provide direct insight into the underlying biology. By linking approaches that visualize brain activity to questionnaires in the context of individualized prediction, we can gain new insights into the biology and behavioral aspects of brain health. Resting-state fMRI (rs-fMRI) can be used to identify biomarkers of these conditions and study patterns of abnormal connectivity. In this work, we estimate mental health quality for individual participants using static functional network connectivity (sFNC) data from rs-fMRI. The deep learning model uses the sFNC data as input to predict four categories of mental health quality and visualize the neural patterns indicative of each group. We used guided gradient class activation maps (guided Grad-CAM) to identify the most discriminative sFNC patterns. The effectiveness of this model was validated using the UK Biobank dataset, in which we showed that our approach outperformed four alternative models by 4-18% accuracy. The proposed model’s performance evaluation yielded a classification accuracy of 76%, 78%, 88%, and 98% for the excellent, good, fair, and poor mental health categories, with poor mental health accuracy being the highest. The findings show distinct sFNC patterns across each group. The patterns associated with excellent mental health consist of the cerebellar-subcortical regions, whereas the most prominent areas in the poor mental health category are in the sensorimotor and visual domains. Thus the combination of rs-fMRI and deep learning opens a promising path for developing a comprehensive framework to evaluate and measure mental health. Moreover, this approach had the potential to guide the development of personalized interventions and enable the monitoring of treatment response. Overall this highlights the crucial role of advanced imaging modalities and deep learning algorithms in advancing our understanding and management of mental health.

## 1 Introduction

Resting-state functional MRI (rs-fMRI) has become one of the most widely used modalities for analyzing functional links to mental health in the human brain. By analyzing differences in brain connectivity patterns using rs-fMRI, researchers can gain insight into the neural substrates of mental health and potentially identify biomarkers for healthy brain function Goulas and Margulies (2021). Studies have demonstrated that the efficacy of rs-fMRI network-based classification can be significantly improved using deep learning techniques Li et al. (2021). These advancements have opened the possibility of applying this classification approach to the fast and objective diagnosis of mental conditions such as major depressive disorder Uyulan et al. (2021), schizophrenia Liu et al. (2022), anxiety disorder Al-Ezzi et al. (2021), bipolar disorder Cheng et al. (2022) and post-traumatic stress disorder Saba et al. (2022). Recent studies use rs-fMRI and deep learning to predict cognitive decline in healthy aging individuals Chen, Liu, Zhu, Gu, and Wang (2021), showing potential for individualized interventions aimed at promoting and maintaining mental health.

The majority of current clinical criteria for determining the severity of mental health symptoms rely on the subjective assessment of the patient’s symptoms and self-reported medical history. More recently, predictive modeling based on machine learning (ML) has been used to interpret neuroimaging data to determine symptom severity or cognitive impairment. The neuroimaging field has shown a rising interest in ML technologies due to challenges in integrating an enormous amount of information in neuroimaging scans. ML algorithms are mathematical models created to identify patterns in known data and use that information to predict patterns in new data. This approach can be applied to clinical populations, including individuals with mental health conditions. For example, functional connectivity (FC) between brain regions measured by rs-fMRI in depressed individuals demonstrate distributed variations across the entire brain Craddock, Holtzheimer III, Hu, and Mayberg (2009). Another study Zeng et al. (2012) utilizing FCs and linear support vector machines (SVM) achieved an accuracy of 94% while classifying between patients with depression and healthy controls. Similarly, a statistical machine learning method such as partial least squares (PLS) regression was used to predict different clinical measures, such as the Positive and Negative Affect Schedule (PANAS), Beck Depression, Inventory-II (BDI-II), Snaith-Hamilton Pleasure Scale (SHAPS), and age from functional connectivity data Yoshida et al. (2017). These predicted clinical scores were further used to classify the depressed patients from healthy controls with 80% accuracy.

Nevertheless, recent advances in deep learning approaches show that, particularly for complex high-dimensional datasets such as fMRI data, the deep models show a significant improvement in performance over standard ML models Su, Xu, Pathak, and Wang (2020). Deep learning algorithms may be trained to recognize abnormalities in fMRI data that are linked with certain mental health problems, and these features can then be used to identify the existence or intensity of a mental health condition. It can also be used to produce tailored treatment options for people with mental health concerns in addition to identifying and predicting them. Furthermore, deep learning models have been successfully utilized on raw fMRI data to perform classification tasks such as detecting distinct brain states or conditions Riaz et al. (2018). Convolutional neural networks (CNN) are amongst the most widely used deep learning models for connectome-based classification and this is particularly significant given how well CNN performs in image classification as well as object recognition Kawahara et al. (2017). Yet, the accessibility of a significant number of training samples is a crucial need for deep learning approaches. Hence, very basic CNN models should be constructed for fMRI-based applications, in accordance with the quantity of data that is accessible.

This paper focuses on classifying participants into different mental health categories based on sFNC data from rs-fMRI. The self-reported behavioral measures of mental health from the UK Biobank were aggregated to obtain a mental health score for each subject. These mental health scores were subjected to Gaussian mixture model (GMM) clustering to obtain the optimum number of categories for classification. Finally, the labels for classification were obtained after placing the participants into four different classes such as excellent, good, fair, and poor mental health. Following this, the sFNC features were input into a one-dimensional convolutional neural network (1D-CNN) to extract useful connectivity parameters for categorizing mental health. The key contributions of this study are as follows: (1) the novel method used a combination of neuroimaging data and a set of self-reported assessment data on questions related to mental health to provide a flexible prediction of mental health quality; (2) the automatic computation of subcategories of mental health quality in any population; (3) interpreting the deep learning model by identifying salient regions in the sFNC associated with each mental health quality category; (4) improved generalization and robustness by training and optimizing the deep learning model on a large dataset; and (5) the model also demonstrated superior performance when compared to other state-of-the-art machine learning algorithms.

## 2 Methods

### 2.1 Participants

The data for this analysis were acquired from the UK Biobank database Miller et al. (2016). The sample comprised 34606 participants, whose ages ranged from 53 to 87 (69.75±7.43) years as shown in Table 1. Participants included 19120 females (53.1%) and 16880 males (46.8%).

**Table 1:**
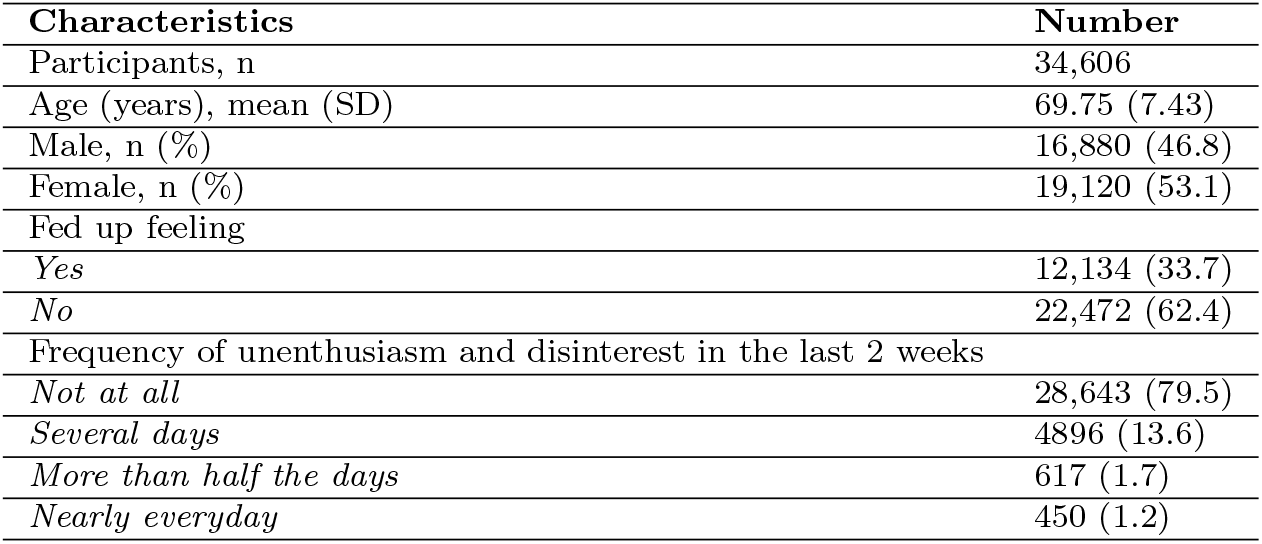
Demographic information from the UKBioank database.

### 2.2 fMRI data acquisition and preprocessing

A 32-channel head coil 3-Tesla (3T) Siemens Skyra scanner was used to scan all the participants. Next, resting-state fMRI images were obtained using a gradient-echo echo planar imaging (GE-EPI) technique. The acquisition parameters consist of no iPAT, fat saturation, flip angle (FA) = 52^*°*^, spatial resolution = 2.4 *×*2.4*×* 2.4mm, field-of-view (FOV) = (88 *×*88*×* 64 matrix), repeat time (TR) = 0.735s, echo time (TE) = 39 ms and 490 volumes. Also, eight slices were acquired concurrently and hence the multiband acceleration factor was set to eight. During the 6min and 10s resting-state scanning phases, participants were asked to passively look at a crosshair and stay relaxed.

We conducted several preprocessing procedures on the UK Biobank database. To reduce the effects of subject-specific motion, we used MCFLIRT Jenkinson, Bannister, Brady, and Smith (2002), an intra-modal motion correction tool. To evaluate brain scans among participants, we employed grand-mean intensity normalization to scale the full 4D dataset by a single multiplicative factor. Further, to eliminate residual temporal drifts, we filtered the data with a high-pass temporal filter and rectified geometric aberrations using FSL’s Topup tool Andersson, Skare, and Ashburner (2003). After EPI unwarping, we employed a gradient distortion correction (GDC) unwarping stage. Next, structural artifacts were eliminated using Independent Component Analysis (ICA) along with FMRIB’s ICA-based X-noiseifier Salimi-Khorshidi et al. (2014). Furthermore, the data were standardized to an MNI EPI template with FLIRT, succeeded by SPM12. Finally, the data were smoothed with a Gaussian filter with a full width at half maximum (FWHM) of 6mm.

Following preprocessing, we applied a completely automated spatially constrained ICA using the NeuroMark Du et al. (2020) technique on the resting state-fMRI data. First, independent components (ICs) were calculated using two large-sample healthy control datasets (HCs). Second, replicable intrinsic connection networks (ICNs) were obtained by comparing and evaluating the spatial maps of ICs from various datasets. The highly replicated ICNs were then used as network templates in an adaptive ICA technique to automatically estimate subject-specific functional networks and related time courses (TCs). Finally, several functional connectivity parameters were computed and assessed; these included static and dynamic functional network connections. According to their functional and anatomical characteristics Allen et al. (2014), these ICNs were classified into seven functional domains: subcortical (SC: 5 ICNs), auditory (AUD: 2 ICNs), sensorimotor (SM: 9 ICNs), visual (VIS: 9 ICNs), cognitive control (CC: 17 ICNs), default mode (DM: 7 ICNs), and cerebellar (CB: 4 ICNs). We used the sFNC as input to our model and as the basis for all subsequent analyses in this study.

### 2.3 Mental health category identification

The self-reported questionnaires from 34606 participants were collected from the UKBiobank database for creating the labels. While the UK Biobank contains mental health data from different sources, we focus on using the assessment center questions completed using the touch screen on the day of the scan. Table 2 shows the 20 questions and the corresponding responses for each question. Here we normalized the responses to the questions so that it ranges from 0 to 1. For instance, in the case of mood swings, ‘0’ corresponds to no mood swings, and ‘1’ corresponds to having mood swings. Whereas in the case of frequency of depressed mood in the last 2 weeks, ‘0’ denotes not at all, ‘0.33’ denotes several days, ‘0.67’ denotes more than half the days, and ‘1’ denotes nearly every day of depressed mood.

**Table 2:**
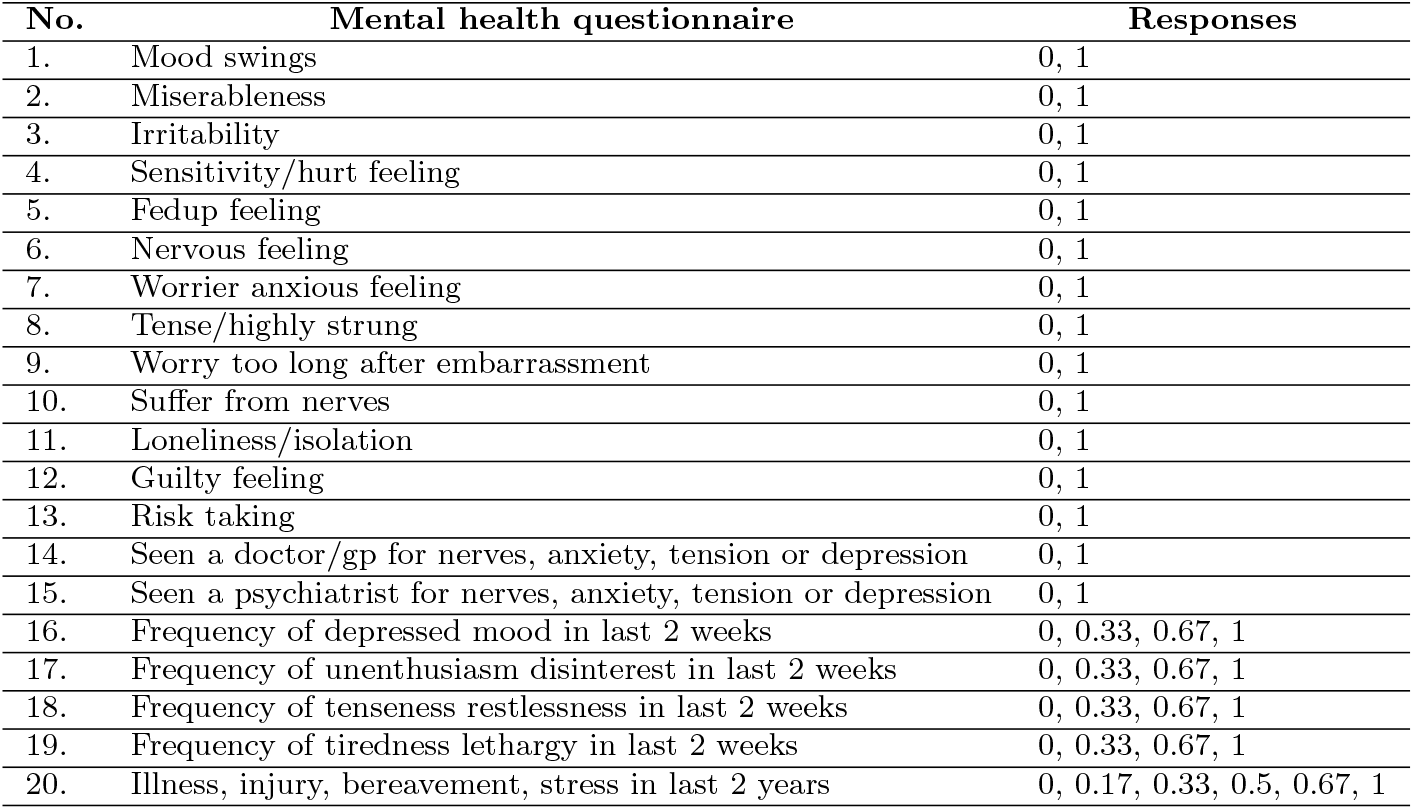
The questions and responses related to mental health that are used in this study from the UKBioank database.

The first 12 questions included in the table enable the calculation of the Eysenck Neuroticism (N-12) score. Individuals with high neuroticism scores are more prone to negative moods and to experience sensations such as anxiety, worry, fear, wrath, frustration, and loneliness. As a result, people with high neuroticism scores are regarded to be at risk of developing mood disorders, anxiety disorders, and substance use disorders Barlow, Curreri, and Woodard (2021). On the other hand, questions 16-19 reflect recent depressive symptoms (RDS-4), a continuous measure of depression symptom severity acquired at the time of scanning. The RDS-4 uses four self-report questions to measure low mood, indifference, restlessness, and weariness. Each question inquires about current symptom occurrences, especially within the past 2 weeks. The four response alternatives are: not at all, several days, more than half the days, and practically every day. In comparison to N-12, RDS-4 assesses the current state of depressed symptoms, whereas N-12 assesses personality traits. Later Smith and colleagues created a categorical (case-control) measure of the lifetime incidence of depression using questions from the evaluation data Smith et al. (2013). This was represented using questions 14 and 15 and they served as an indication of the subject’s probable depressive status. Nevertheless, these questions did not distinguish between isolated and recurring depressive episodes. For instance, if the participants indicated they had seen a doctor or a psychiatrist for nerves, worry, stress, or depression, their depression status was set to 1.

In this study, we selected 20 questions for each subject based on previous studies conducted in UK Biobank associated with mental health Dutt et al. (2022). A mental health score was calculated for each subject by summing their normalized responses to the questions. Here the maximum possible score for any subject is 20 and the minimum score is 0. A histogram of the mental scores over the 34606 participants is shown in Fig. 1. According to the histogram, the primary conclusion is that participants with low mental health scores have excellent mental health while those who score closer to 20 have poorer mental health. In this problem, the number of categories of mental health quality is not predefined. Hence we use the Gaussian Mixture Model clustering Fraley and Raftery (2002) method to automatically find the different groups present in the data.

**Fig. 1:**
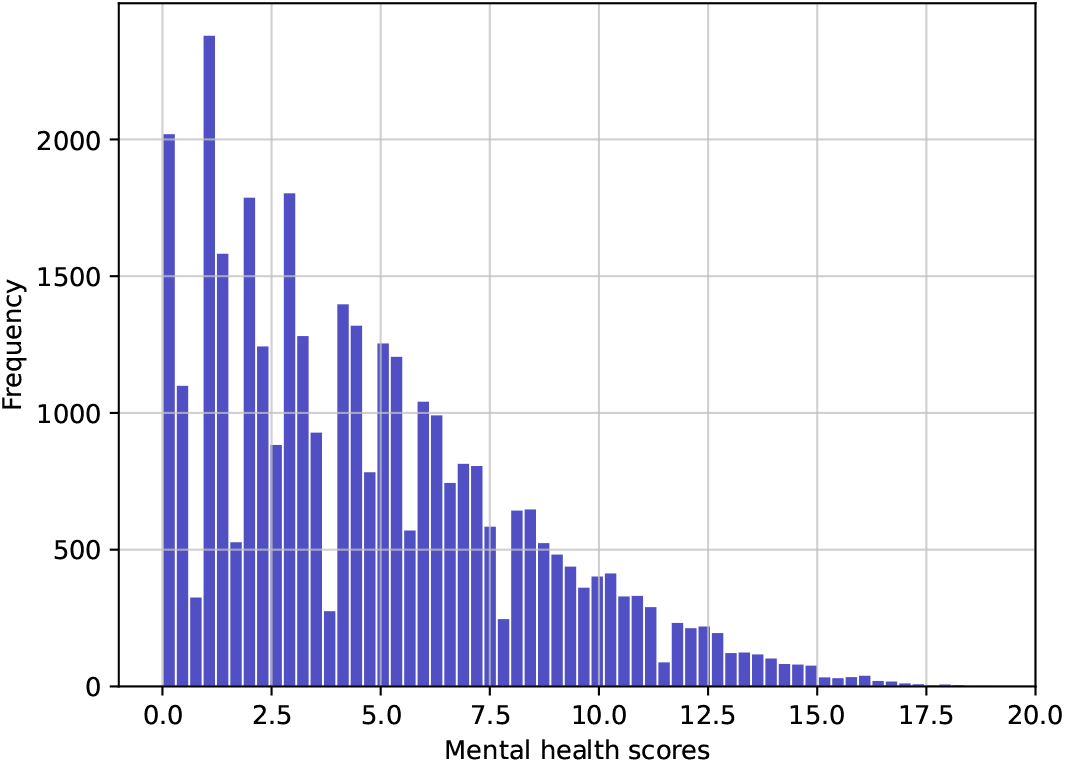
Histogram of the mental health scores for 36,000 participants from the UKBiobank database.

GMMs are unsupervised probabilistic models that follow the assumption that all data points are generated from a fixed set of Gaussian distributions.

This approach distributes data points into distinct groups using the soft clustering technique. Multiple Gaussian distributions are fitted to the data and the distribution parameters such as mean, variance, and weight are calculated for each cluster. The probability of each data point belonging to a cluster is determined after learning these parameters. Given the one-dimensional mental health scores data, the probability density function is computed as follows:

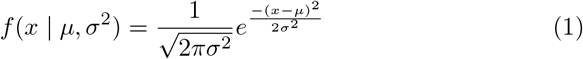

Here *x, µ*, and *σ*^2^ represent the data points, mean, and variance respectively. Expectation maximization Dempster, Laird, and Rubin (1977) is used to estimate the mixture model’s parameters when the number of clusters is known. This is an iterative strategy with the property that the maximum likelihood of the data strictly rises with each additional iteration. There are two phases in the expectation-maximization process. Initially, the mean and variances are assigned randomly. Next, the posterior probability that each data point belongs to a cluster is determined in the expectation phase using the current mean and variances. The cluster means and variances are recalculated in the maximization stage using the probability obtained in the expectation step. The steps are repeated to get a maximum likelihood estimate until the convergence of the algorithm.

We used Bayesian Information Criterion (BIC) Schwarz (1978) to determine the ideal number of clusters. The BIC compares the maximum likelihood function with the number of model parameters, k, and data points, n. Adding a penalty for the number of model parameters allows it to choose the model with the fewest parameters that best describe the data. Thus the model with the lowest BIC score is selected from a finite set of models. The BIC score is calculated using the below equation:

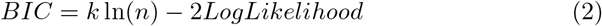

Analysis of the mental health scores resulted in four clusters. The labeling of the clusters was as follows: excellent (score range = 0 to 3), good (score range = 4 to 7), fair (score range = 8 to 11), and poor(score range = 12 to 20). We calculated the range of the mental health quality groups using the mean and standard deviation of their associated clusters. The maximum value of the range is given by adding the mean with twice the standard deviation, whereas the minimum value of the range is obtained by subtracting twice of standard deviation from the mean.

### 2.4 Predictive modeling using functional network connectivity data

The prediction network for the sFNC data is designed using a one-dimensional convolutional neural network (1D-CNN) Krizhevsky, Sutskever, and Hinton (2017); LeCun, Bengio, and Hinton (2015). Our main goal was to predict mental health quality categories using the sFNC data from the UK Biobank database. The features consist of 53 *×* 53 sFNC matrices that represent interconnectivity strengths between various ICNs. The corresponding labels are the mental health scores that we computed from the self-report data.

The proposed CNN is a 16-layer network with four convolutional layers with a kernel size of three and with 16, 32, 64, and 128 filters respectively. The rectified linear unit (ReLU) Nair and Hinton (2010) non-linearity is used in the convolutional layers. The three fully connected layers towards the end of the model have 64, 16, and 4 nodes and on the output layer, the softmax activation function Bridle (1990) is used to determine the likelihood that each sample belongs to a class. The model uses four max-pooling layers with kernel size two to decrease the dimensionality of the feature maps and limit overfitting. Also, a drop-out regularization Srivastava, Hinton, Krizhevsky, Sutskever, and Salakhutdinov (2014) with a probability of 0.2 and 4 layers of batch normalization was added for regularization. The Adam optimizer Kingma and Ba (2014) was used to train the proposed CNN for 150 iterations, with a learning rate of 0.001 and a batch size of eight. The data set underwent five-fold cross-validation in which four folds are used for training and the remaining fold was used for testing. The excellent, good, fair, and poor categories consisted of 16256, 11845, 5058, and 1447 participants respectively. Since the data for the four classes was imbalanced, Synthetic Minority Over-sampling Method (SMOTE) Chawla, Bowyer, Hall, and Kegelmeyer (2002) was used to construct a balanced dataset. This is a method of oversampling the minority class by producing synthetic samples rather than oversampling using replicated actual data values. In accordance with the amount of over-sampling needed, neighbors from the k nearest neighbors of a minority class are picked at random. Fig 2 (C) illustrates the architecture of the proposed model.

**Fig. 2:**
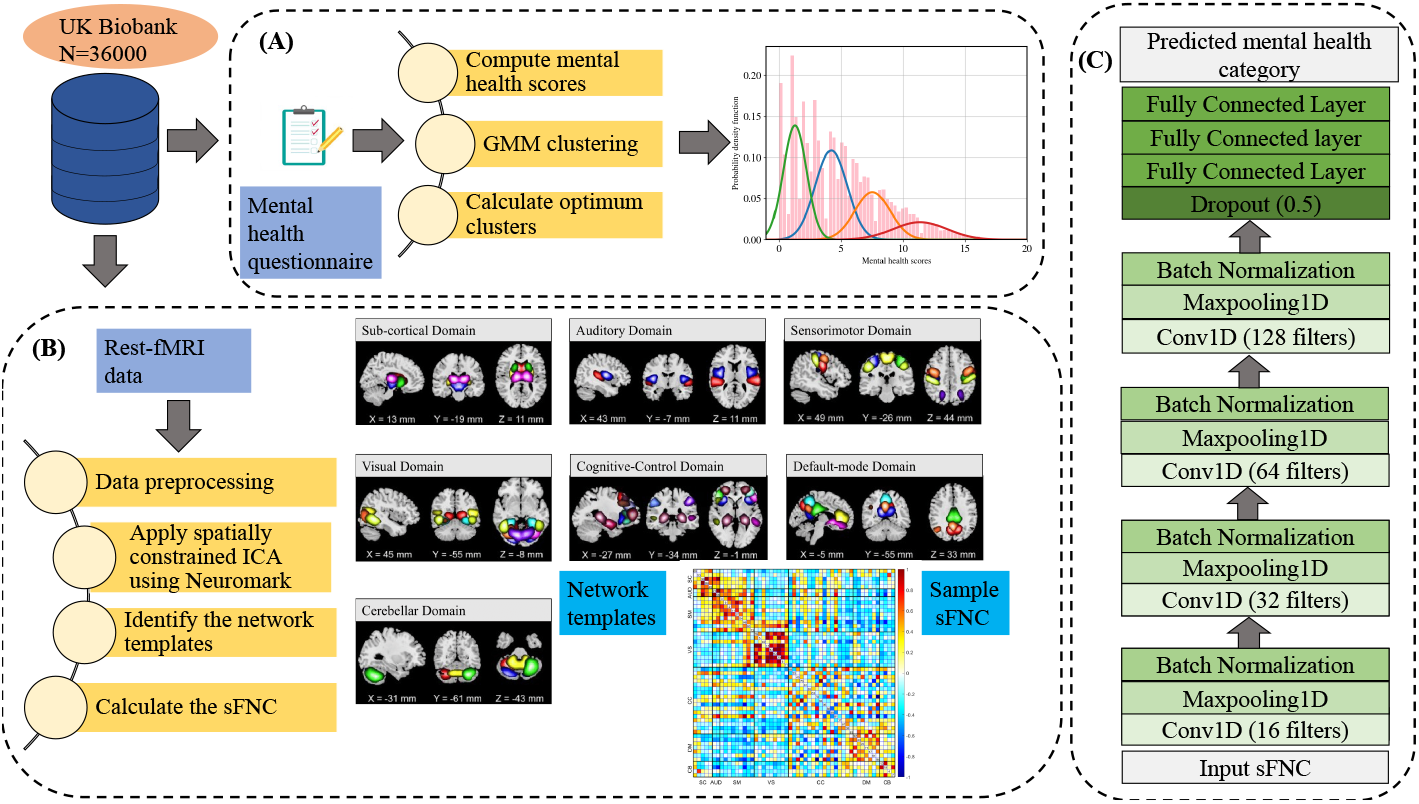
The proposed deep learning architecture for mental health category prediction.

## 3 Results

### 3.1 Performance evaluation using various metrics

We used performance measures such as sensitivity, specificity, precision, and accuracy to analyze the multiclass classification. As illustrated in Fig.3, the poor category had the highest values (all *>* 95%) for all performance measures. Also, for all participant groups, specificity was higher than sensitivity. This indicates that there are fewer false positives than false negatives present in the test set after classification. Moreover, the excellent and the good categories have lower precision values than the fair and poor categories. Hence when the proposed model was inaccurate, incorrect classifications were more likely to occur between these two categories. Additionally, for all categories, accuracy exceeded 75%.

**Fig. 3:**
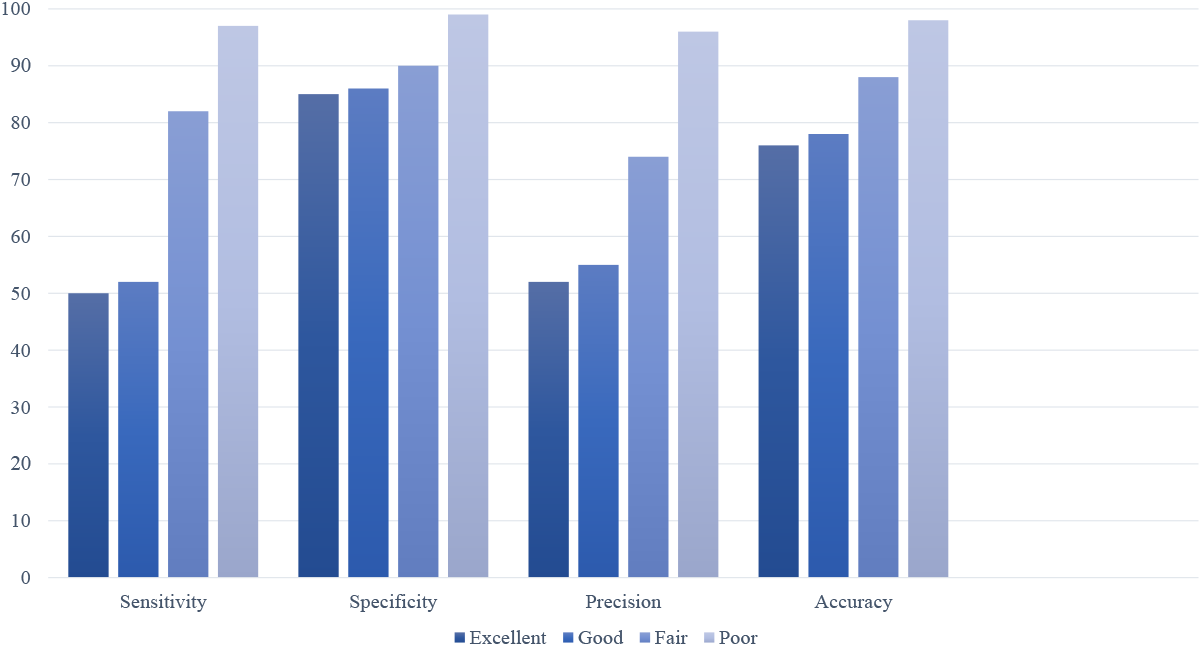
Comparison of performance metrics for the mental health categories using the proposed method.

The Receiver Operating Characteristic (ROC) curves of the proposed model were examined to measure its effectiveness in performance. The capacity of the model to discriminate between participants belonging to various groups according to different thresholds is shown by the area under ROC curves (AUC). Higher AUC values indicate that the model is more accurately classifying the participants. Fig 4 displays the multiclass ROC curve for the 1D-CNN for the four classes of mental health. The closer the curves are to the top-left corner, the better the performance of the model. We used the one vs. all methods to depict the four ROC curves for the different categories in this multiclass model. The proposed model yielded AUCs of 0.81, 0.81, 0.93, and 1 for the excellent, good, fair, and poor mental health categories respectively. Table 3 shows the accuracy for each class and the total average accuracy for the different classifiers to evaluate the classification performance. We compared the performance of the proposed network with four other baseline models (support vector machines (SVM), multi-layer perceptron (MLP), random forests (RF), and naïve Bayes classifier (NB)). The hyperparameter tuning for these classifiers was completed using the grid search method. We can see that the proposed CNN clearly outperformed all the conventional machine learning classifiers in distinguishing among the mental health categories. It achieved the best accuracy of 76%, 78%, 88%, and 98% for excellent, good, fair, and poor respectively and it had an overall average accuracy of 85%. Also, the proposed model improved the average accuracy by 5%, 9%, 4%, and 18% over SVM, MLP, RF, and NB, respectively. This indicated the effectiveness of learning complex, deep characteristics from fMRI data.

**Table 3:**
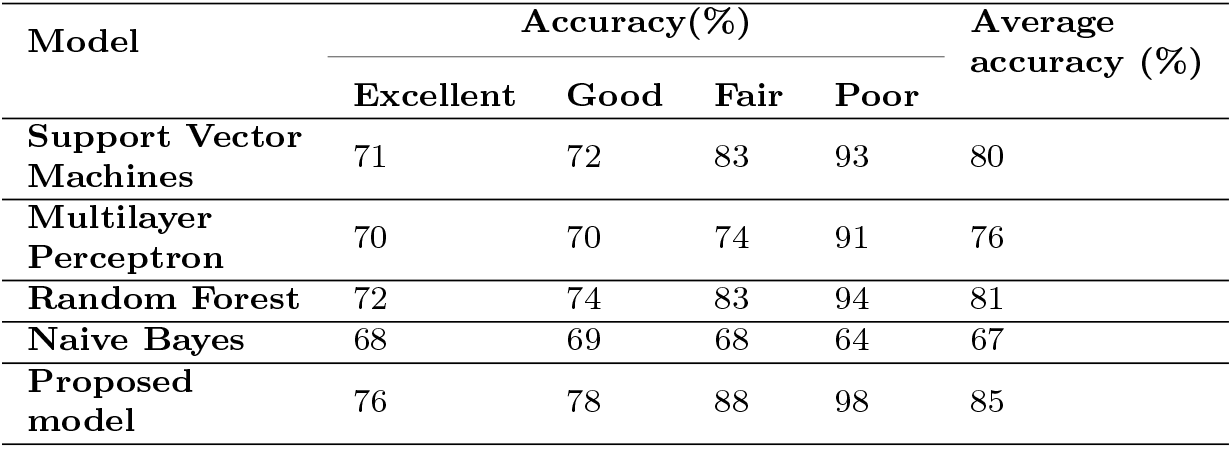
Accuracy comparison between the proposed model and the state-ofthe-art techniques.

| Model | Accuracy(%) |  |  |  | Average accuracy (%) |
| --- | --- | --- | --- | --- | --- |
|  | Excellent | Good | Fair | Poor |  |
| Support Vector Machines | 71 | 72 | 83 | 93 | 80 |
| Multilayer Perceptron | 70 | 70 | 74 | 91 | 76 |
| Random Forest | 72 | 74 | 83 | 94 | 81 |
| Naive Bayes | 68 | 69 | 68 | 64 | 67 |
| Proposed model | 76 | 78 | 88 | 98 | 85 |

**Fig. 4:**
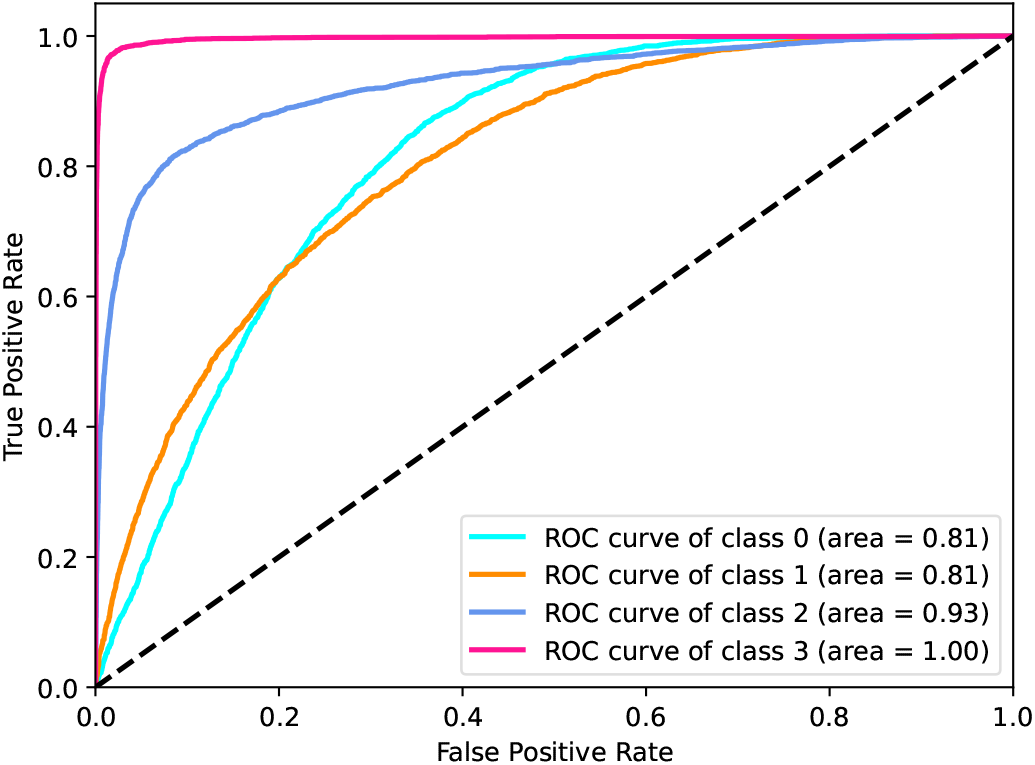
ROC curve for different mental health categories. Here class 0 denotes the excellent category, class 1 is good, class 2 is fair and class 3 is the poor category.

### 3.2 Analysis of the mental health questionnaires

Responses to 20 mental health questions were analyzed to identify which questions best discriminated the participants in each of the mental health quality categories. We calculated the percentage of scores within each category by summing the scores of the questions for the participants assigned to each category, dividing it by the total number of participants, and multiplying it by

100. Here the higher percentage value denotes that the corresponding question contributed more to the specified category. Fig. 5 presents a bar graph representing these percentage values for each mental health category. For the excellent category, all questions had very low percentage values (less than 30%). Here we observe that extremely low values can be found for symptoms such as nervous feelings, tense feelings, suffering from nerves, loneliness/isolation, frequency of depressed mood, unenthusiasm, and restlessness in the last 2 weeks and for seeing a psychiatrist for nerves, anxiety, and depression. The lack of these symptoms can be considered an indication of excellent mental health quality. When a person is not experiencing these negative emotions, it may signify that they are able to effectively regulate their emotions, which can further improve their general well-being. In contrast, for the poor category, 14 questions had more than 50% value. Hence poor mental health directly corresponds to symptoms such as mood swings, miserableness, irritability, sensitivity/hurt feelings, fed-up feelings, nervous feelings, worry/anxious feelings, tense feelings, worrying too long, suffering from nerves, loneliness/isolation, guilty feelings, frequency of tiredness and lethargy, and seeing a doctor for nerves, anxiety, and depression. The fair category shows similar patterns but with low percentage levels.

**Fig. 5:**
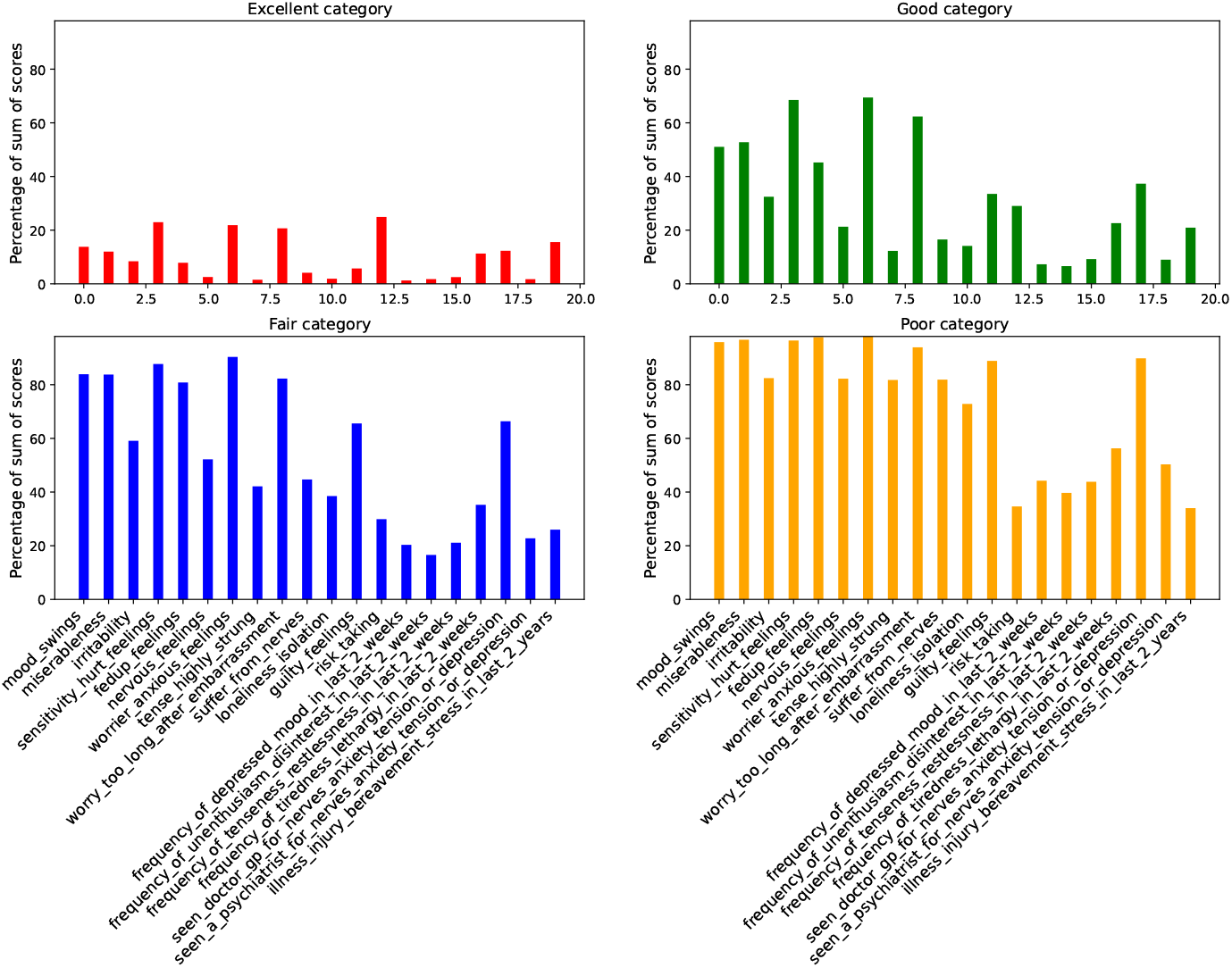
Contribution of each question towards the mental health category.

### 3.3 Mean sFNC and connectogram analysis

In our study, we calculated mean sFNC for participants in order to assess the neural connectivity patterns associated with each of four empirically derived mental health quality categories. In Fig. 6 for the excellent category, a sparsely weakly connected sFNC was present in the VS-SM and CC-SM domain pairs. In the case of the good category, the connectivity was highly present in the SMSC and VS-SC domain pairs. For the fair group, however, richer connectivity patterns were apparent in the SC domains and VS-SM domain pairs. Many domains showed high connectivity values at random places as we approached the poor category, but a large cluster could be seen in the SM and VS-SM domain pairs.

**Fig. 6:**
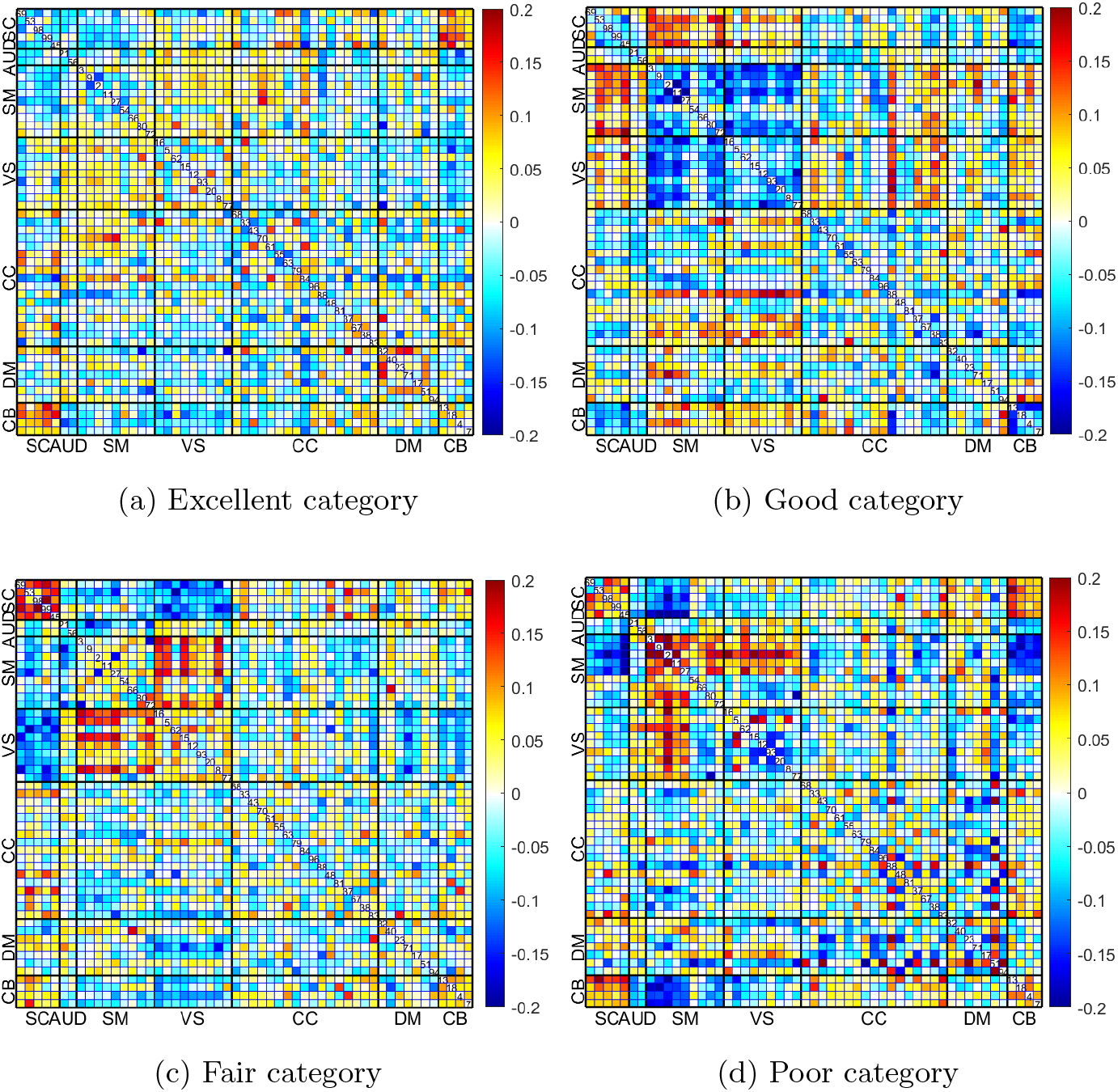
Mean sFNC of the different test participants for the various categories.

A connectogram analysis is done in Fig. 7 (a) and (b) to illustrate the differences between the excellent and poor mental health quality categories. In this plot, the blue color denotes a negative correlation, while yellow denotes substantial links with a positive association. The degree of relevance is indicated by the opacity of the lines. The connectogram for the excellent category exhibited significantly negative values in two connection pairs, including the DM-VIS, and DM-SM domains. In comparison with the excellent group, the poor category showed a positive association in the DM and SM domain interactions. Moreover, as we visualize the progression of the connectivity pattern from excellent to poor mental health category in Fig. 7 (c), certain ICs in the SM, VIS, and CC domains show a monotonic increase in interactions.

**Fig. 7:**
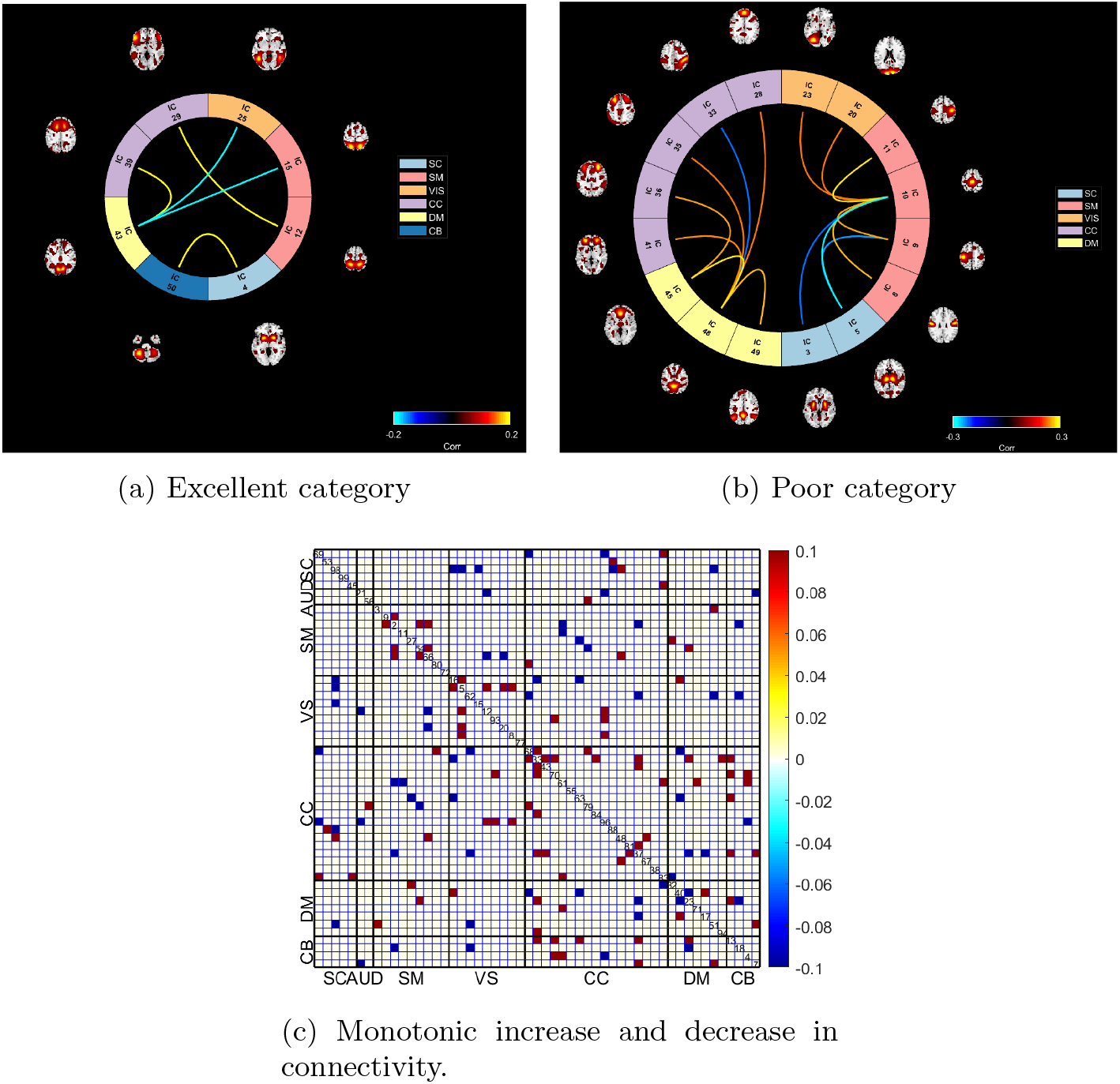
(a) Connectogram of the mean sFNCs of participants belonging to the excellent category. (b) Connectogram of the mean sFNCs of participants belonging to the poor category. In both cases, warmer colors represent increased connectivity and cooler colors denote decreased connectivity. (c) The red and blue colors denote a monotonic increase and a monotonic decrease in connectivity in the sFNCs.

### 3.4 Saliency maps of the sFNC

Interpretable models are critical for improving predictability and increasing clinical acceptability. We employed guided gradient class activation maps (guided Grad-CAM) Selvaraju et al. (2017) to objectively define the prominent areas of the sFNC contributing to mental health category prediction in order to generate interpretable results for the network predictions. Then, we computed the gradients for every prediction score with respect to the extracted feature maps from the final convolution layer. The weights for feature significance were then calculated using the global average pooling of these gradients. ReLU is applied after a weighted mixture of forward activation maps. ReLU has the advantage of highlighting features that have a positive influence on the target class. It has been shown that localization maps without ReLU may contain more information than the target class, such as negative pixels that likely correspond to other categories Zhou, Khosla, Lapedriza, Oliva, and Torralba (2016). While the target class-specific localization map resulting from this might be significant, it could end in the loss of key context and global information. Fig 8 illustrates the saliency maps for the four mental health categories. The localization maps were averaged across all participants to create the saliency maps. The regions that the model considers salient (darker red zones) for the prediction of mental health are highlighted by overlaying the saliency map from guided Grad-CAM. The disadvantages of employing the guided Grad-CAM technique include the inability to generalize this approach across larger samples where patient saliencies and varied average sFNCs might lead to ambiguous activations. In the case of misleading activations, the saliency map highlights regions that are not actually relevant to the target class. This further leads to the calculation of average class activation maps containing missing relevant information and thereby providing limited inferences. Moreover, the current interpretability approaches relying on class activations are tailored to individual participants.

**Fig. 8:**
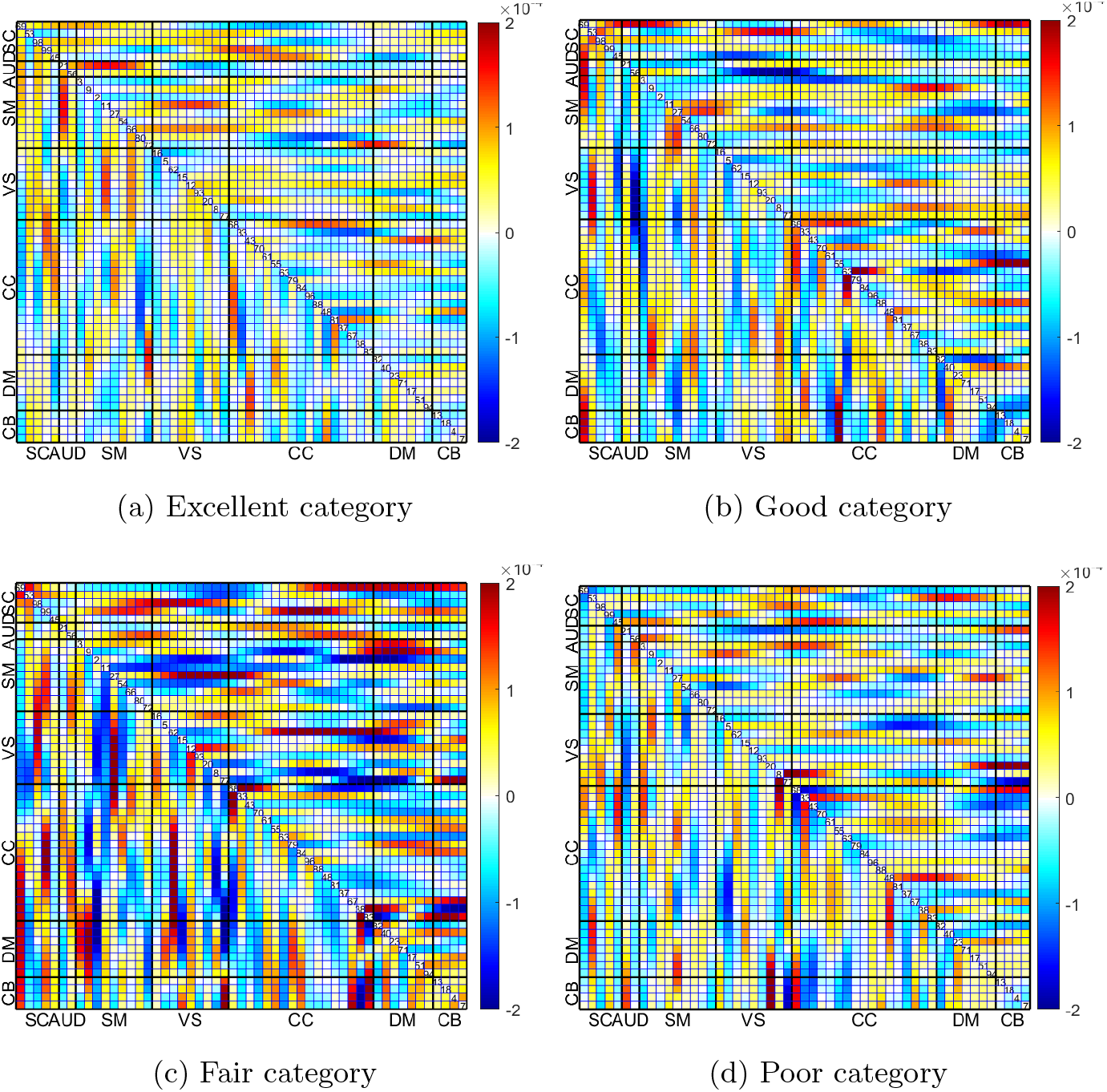
Saliency maps of different test participants for the various categories.

## 4 Discussion

The aim of this study was to develop a framework for sFNC analysis to provide insights into the relationship between self-reported mental health assessment data and sFNCs that comprise the different brain regions. Using a deep learning technique, we discovered four distinct categories of mental health, characterized these groups based on various self-report patterns, and found discriminative functional connectivity areas. The mental health questionnaire analysis also revealed which questions best-distinguished individuals in each of the mental health quality categories. Importantly, all these findings were achieved on a large dataset which further increased the robustness and generalization of the deep learning method.

Recent studies have suggested that depression, anxiety, stress, and other mental health conditions may be associated with disparities of interconnections among brain regions rather than increased or decreased activity of individual areas Wang et al. (2019); J.-T. Zhang et al. (2016). Hence, researchers studying mental health have shifted the focus of imaging studies to connections among brain areas. The data-driven connectome-based predictive models take brain connectivity data as input and generate predictions of behavioral measures in participants Shen et al. (2017). Here the predictive model assumes a linear relationship between the connectivity data and the behavioral variable. But these models may not be optimal for capturing complex, nonlinear relationships between connectivity and behavior. Deep learning models, on the other hand, can learn complicated non-linear correlations between variables by using nonlinear activation layers. Hence in our study, we used the one-dimensional convolutional neural networks to enhance the prediction accuracy by capturing the patterns present between the sFNCs and the self-reported assessment data. Moreover, deep learning models had shown promise in sFNC-based predictive modeling in previous research, but their lack of interpretability has remained a concern Cwiek et al. (2022). To address this issue, we used saliency maps to highlight the most significant regions of the sFNC matrix that contribute to the model’s prediction. This served as a powerful tool for the interpretability of the deep learning model.

Using the connectogram analysis, we also visualized and identified relevant brain connections contributing to the prediction model. Various studies examining stress-induced neural responses and recovery patterns through post-stress rs-fMRI scans show that the overall intra- and inter-network FC of certain core networks have been frequently reported to be altered after acute stress. These regions consist of the default mode network (DMN) Clemens et al. (2017) which is involved in internally-directed cognition and includes the posterior cingulate cortex (PCC). Also, the salience network (SN) W. Zhang et al. (2019); X. Zhang, Huettel, O’Dhaniel, Guo, and Wang (2019) includes the anterior insula and the dorsal anterior cingulate cortex, which detects behaviorally relevant stimuli and reallocates the brain’s neural resources. Relative to the findings in these previous studies, we also found that in the case of the poor mental health category with an increased likelihood of symptoms of stress, anxiety, and depression, there was also an increase in connectivity in the default mode and sensorimotor regions as shown in the connectogram. Overall, we demonstrated that the proposed architecture significantly improves the accuracy with which one can classify individual participants into distinct mental health quality categories. The saliency analysis also provides several sFNC pairs that exhibit a strong connection in predicting a subject’s mental health. In the case of the excellent category, the significant sFNC regions are AM-AUD, VS-SM, SM-DM, and CC-DM. The sFNC is sparsely distributed and consistent with the mean sFNC plot shown in Fig. 6 (a). The good category on the other hand has dominant regions in SM, CC, and SC domains. While the fair category has a mixture of both positive and negative significant regions with the CB domain being one of the clearly contributing regions of this category. Finally, in the case of the poor category, the DM-VS, CC, and DM represent the regions that have the most positive influence in this category. As a result, deep learning is a promising strategy for assisting healthcare providers in the development of neuroimaging-based biomarkers for earlier detection in clinical settings. The research conducted may also be employed to help in monitoring mental health quality and response to interventions. Additionally, the findings can serve as a guide for diagnostic testing and therapies that aim to enhance the participants’ quality of life.

## 5 Conclusion

In this paper, we developed a deep learning framework based on 1D-CNN for categorizing mental health scores. On a large dataset, this model proved more effective than conventional machine learning techniques in classifying individual people accurately. In fact, this model has several advantages as it uses both sFNC and self-reported assessment data for predictive analysis. First, sFNC provides objective measurements of brain activity and connectivity, whereas using just the self-reported data to obtain mental health quality can be biased or influenced by social desirability Abdallah, Farrugia, Chirokoff, and Chanraud (2020). Secondly, sFNC data provides information about functional networks across the whole brain, whereas self-reported data may only capture information about specific symptoms. Combining this information through deep learning provides a more comprehensive view of brain function and connectivity. This can help to identify patterns of connectivity that are associated with specific symptoms associated with mental health quality. Finally, the proposed model can clearly distinguish between the different mental health quality groups with high accuracy. Once this has been trained on a dataset, it can be used to make predictions on new data without adding any additional self-reported measures. A limitation of this research is that it focuses on middle-aged and older persons, and the study includes only twenty self-reported assessments performed on the day of scanning.

Nevertheless, our promising findings offer the possibility that neuroimaging data can be leveraged to facilitate a more accurate categorization of people according to their mental health. The categories that self-reported data yielded had distinctive patterns of connectivity. Future work will focus on the development of a deep learning-based fusion model to forecast brain health by using time courses, sFNCs, and spatial maps. Also, we will evaluate whether a predictive model from neuroimaging data can outperform a predictive model based on assessment data. Additionally, we will extend our model to younger adults and include more self-reported measures that are taken at a time point that was independent of the scan date.

## Author contributions

Meenu Ajith: Conceptualization, Methodology, Manuscript Writing & Editing. Dawn M. Aycock: Manuscript Writing & Editing. Erin B. Tone: Manuscript Writing & Editing. Jingyu Liu: Manuscript Writing & Editing. Maria B. Misiura: Manuscript Writing & Editing. Rebecca Ellis: Manuscript Writing & Editing. Tricia Zawacki King: Manuscript Writing & Editing. Vonetta M. Dotson: Manuscript Writing & Editing. Vince Calhoun: Methodology, Funding acquisition, Manuscript Writing & Editing.

## Data Availability

All data is publicly available at the UK Biobank online database (https://www.ukbiobank.ac.uk/).

## Declarations

### Ethical approval

This research study was conducted retrospectively using human subject data made available in open access by the UK Biobank. Ethical approval was not required as confirmed by the license attached with the openaccess data.

### Consent to participate

Informed consent was obtained from all participants.

### Consent to Publish

All authors have consented to publish.

